# Improved *in situ* sequencing for high-resolution targeted spatial transcriptomic analysis in tissue sections

**DOI:** 10.1101/2022.10.16.512401

**Authors:** Xinbin Tang, Jiayu Chen, Xinya Zhang, Xuzhu Liu, Zhaoxiang Xie, Kaipeng Wei, Jianlong Qiu, Weiyan Ma, Chen Lin, Rongqin Ke

## Abstract

Spatial transcriptomics enables the study of localization-indexed gene expression activity in tissues, providing the transcriptional landscape that in turn indicates the potential regulatory networks of gene expression. *In situ* sequencing (ISS) is a targeted spatial transcriptomic technique, which is based on padlock probe and rolling circle amplification combined with next-generation sequencing chemistry, for highly multiplexed *in situ* gene expression profiling. Here, we present improved *in situ* sequencing (IISS) that exploits a new probing and barcoding approach, combined with advanced image analysis pipeline for high-resolution targeted spatial gene expression profiling. We developed an improved combinatorial probe anchor ligation chemistry using a 2-base encoding strategy for barcode interrogation. The new encoding strategy resulted in higher signal intensity as well as improved specificity for *in situ* sequencing, while maintaining a streamlined analysis pipeline for targeted spatial transcriptomics. We showed that IISS can be applied to both fresh frozen tissue and formalin-fixed paraffin embedded tissue sections for single cell level spatial gene expression analysis, based on which the developmental trajectory and cell-cell communication networks can also be constructed. In conclusion, our method is a versatile molecular tool for targeted spatial transcriptomic analysis.

## INTRODUCTION

Spatial transcriptomics enables us to study gene expression from a new perspective by providing the localization information of detected genes. Previously, spatial gene expression profiling can be achieved by the assistance of methods such as laser-capture microdissection (LCM) that allows to analyze regions of interest in tissue samples ^1^. However, it can’t provide an overview of the transcriptional atlas of the target region and its surrounding regions, thus may lead to missing of information on how genes synergistically expressed in their native context. Currently, the mainstreamed spatial transcriptomic technologies are often referred to methods that can offer localization-indexed gene expression information in a defined area of tissue samples. These methods can be categorized into two approaches, one is based on sequencing of spatially-indexed transcripts and the other is based on imaging *in situ* ^2^. The first approach relies on mapping back the detected transcripts to the original compartmentalized regions in tissue by decoding the barcoded cDNA primers after next-generation sequencing (NGS) ^3^. The second approach includes a series of *in situ* hybridization (ISH) ^4^ or *in situ* sequencing (ISS) ^5^ methods that depend on cyclic imaging to decode the barcodes for individual transcripts directly in their original places, achieving highly multiplexed *in situ* RNA detection by using combinatorial color-coding strategies to overcome the limitation of spectrally-distinguished labels for *in situ* hybridization assays. The former sequencing approach has a high coverage of genes that includes almost all expressed mRNAs but at a lower spatial resolution due to each compartmentalized spot normally contains more than one cells ^6^. The imaging-based approach includes targeted spatial transcriptomic methods that normally analyze a panel of pre-selected genes at a much higher resolution because RNA molecules are detected at single molecule resolution ^7^.

*In situ* sequencing is a type of imaging-based targeted spatial transcriptomic technology. In ISS, rolling circle amplification (RCA) is used to amplify circularized probes designed for specific genes or cDNA generated by *in situ* reversed transcription ^5,8,9^, forming rolling circle amplification products (RCPs) that can be sequenced by NGS chemistry to identify the detected genes based on their barcodes or mapping the sequences to the reference genome ^10^. The classical ISS is based on barcoded padlock probe and RCA for signal amplification combined with the combinatorial probe anchor ligation chemistry (cPAL) ^11^ for decoding the amplification products. As a popular and proven versatile targeted spatial transcriptomic method, ISS has been applied for region-specific gene expression profiling ^12^, *in situ* cell typing as well as novel histopathological discovery ^13,14^, etc. Here, we present improved *in situ* sequencing (IISS) based on the probe designing principle from amplification-based single molecule fluorescent *in situ* hybridization (asmFISH) ^15^ combined with an improved combinatorial probe anchor ligation (icPAL) chemistry for RCA-based *in situ* sequencing library preparation. Specifically, this combinatorial 2-base barcoding strategy was shown to greatly improve the sequencing quality with higher signal intensity and better specificity. More importantly, we proposed a new cell segmentation pipeline based on Cellpose ^16^, a deep learning algorithm, to define individual cell regions *in situ*, so as to generate a spatial gene expression matrix. We showed that IISS can be applied to perform *in situ* gene expression profiling and cell typing in fresh frozen tissue section and more excitingly in formalin-fixed paraffin embedded tissue section. What’s more, based on the spatial gene expression atlas generated by IISS, we were able to construct the carcinoma invasive trajectory and investigate cell-to-cell communications between malignancy and normal cells of the analyzed colon cancer tissue. Thus, we have demonstrated that our IISS technology can be used for precise *in situ* cell typing, showing its great potential for high-resolution targeted spatial transcriptomic analyses.

## RESULTS

### Workflow and implementation of the DLP-based IISS

Based on previously established asmFISH ^15^ and ISS ^5^ method, DLPs are designed to target RNA molecules *in situ*. A DLP pair is constituted by an upstream probe and a downstream probe. Each probe contains a 16 nt target recognition sequence, a 9 nt interrogation probe sequence, a 10 nt anchor primer hybridization sequence and a 10 nt unrelated sequence, resulting in a 45 nt long DLP (Supplementary Figure S1a). The gene-specific barcode is divided into two motifs and placed in each of the 9 nt detection probe hybridization motif. Upon recognition and hybridization on the correct target, the upstream and downstream DLP probes can be ligated, and then circularized by a second ligation mediated by a splint oligonucleotide template. The resultant circular DNA molecules can thereafter be amplified by RCA, generating rolling circle amplification product (RCP) that contains hundreds of repeated complementary sequences of the original DNA circles. These RCPs can then be decoded by sequencing the gene-specific barcodes to reveal their identities (Figure 1a). Based on the original combinatorial probe anchor ligation (cPAL) ^5^ NGS chemistry used for DNA nanoball sequencing and the modified cPAL (mcPAL) ^14^ used for previous ISS, we developed an improved cPAL (icPAL) chemistry for improved *in situ* sequencing (IISS). In icPAL, the interrogation probe contains five fixed bases, two degenerate bases and a barcode constituted by predetermined 2-base encoding motifs (Supplementary Figure S1b).

**Figure 1.**
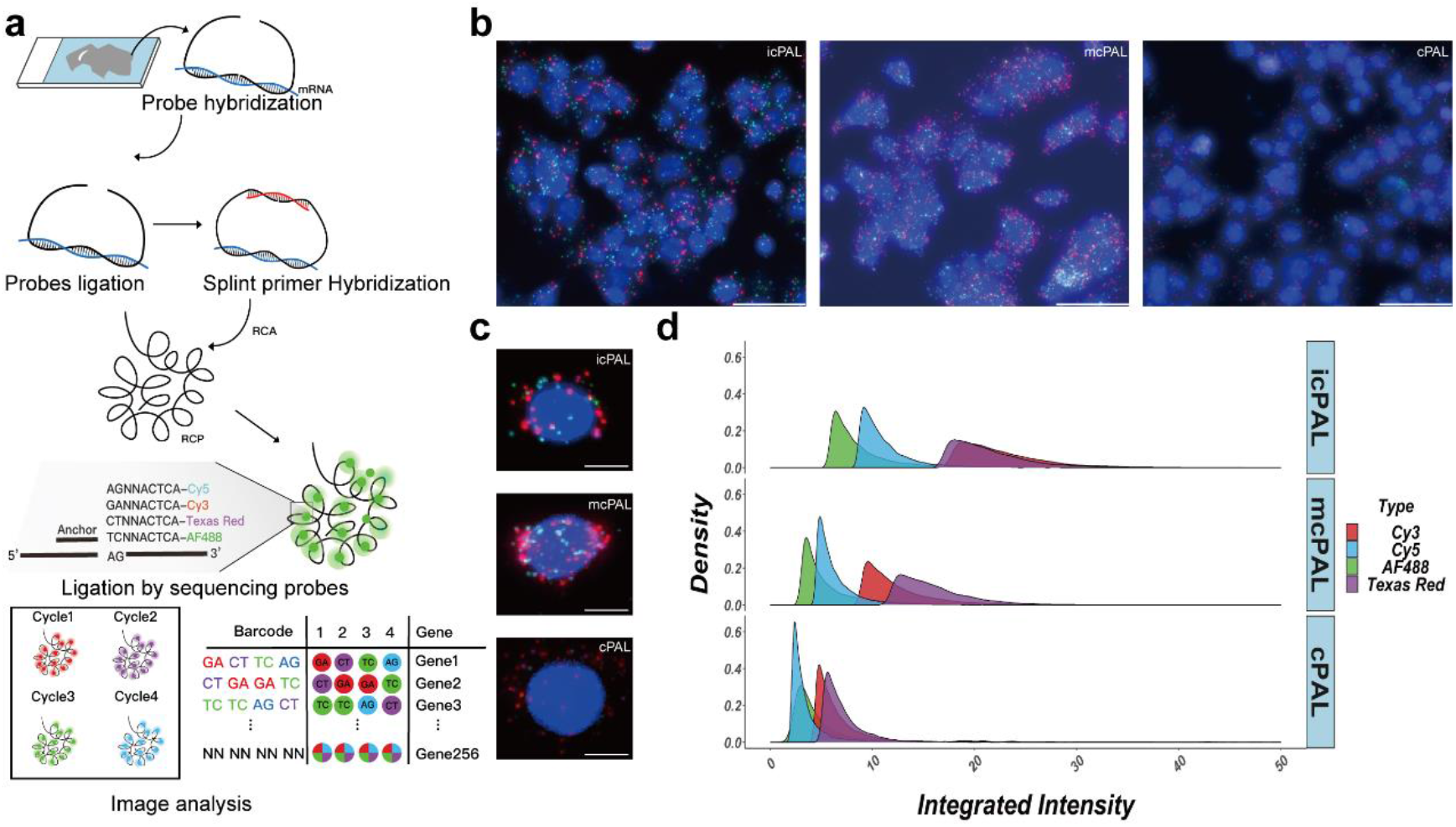
Schematic overview of IISS and the comparison of icPAL with mcPAL and cPAL. (a) Overview of IISS. Gene-specific DLPs hybridize and ligate on the mRNA template, followed by circularization by DNA ligation mediated by a splint oligonucleotide template. Probes that are circularized on the transcripts successfully are then amplified by RCA. Four kinds of sequencing interrogation probes are labelled with unique fluorophores. The barcodes are decoded by image analysis based on identification of the dominant signal. (b) Representative images in Cycle 4 of the sequencing-by-ligation performed on MCF7 slides by icPAL-, mcPAL-, and cPAL-based chemistries. Each channel has been scaled to the same intensity range. Scale bar: 50 μm. (c) Single cell images with DAPI stained nuclei and RCPs of different colors. Scale bar: 10 μm. (d) Comparison of integrated intensities of icPAL, mcPAL, and cPAL in different fluorescence channels.

To compare the performance of cPAL, mcPAL and icPAL, we performed IISS for 20 genes on fixed MCF-7 cell samples. RCPs detected by sequencing-by-ligation chemistry are discrete fluorescence dots of different colors corresponding to the interrogation position in the barcodes (Figure 1b). Enlarged views of single cell images show that icPAL generated the clearest fluorescent signal, in contrast to the blurry dots of mcPAL and lower fluorescent intensity of the original cPAL (Figure 1c). Comparing the integrated intensity of the top 10% RCPs per channel, there was an increase for icPAL, with relative intensities of 15.5 ± 7.5 for icPAL v.s. 9.5 ± 5.33 and 6.1 ± 5.02 (Mean ± SD) for mcPAL and cPAL respectively (Figure 1d and Supplementary Figure S1c). This result thus demonstrates that the 2-base barcoding strategy helped to increase the hybridization and ligation efficiency of the anchor primers and the interrogation probes.

To establish a robust experimental protocol, we then optimized the experimental conditions. Firstly, we compared one-step and two-step sequencing protocols. In the one-step protocol, anchor primers, interrogation probes and DNA ligase were mixed in one-pot and directly incubated with the samples. While in the two-step protocol, the anchor primers were first hybridized with the RCPs, followed by ligation of the sequencing interrogation probes in an independent step. The results showed that there was no obvious difference between one-step and two-step sequencing protocols in terms of the integrated intensity of the top 10% of RCPs (Supplementary Figure S1d). Secondly, we compared the impact of the lengths of the anchor primers. The integrated fluorescence intensity in Cycle 1 and Cycle 2 is higher for 15 nt anchor primer (16.6 ± 6.2) than 11 nt (13.1 ± 5.2), 13 nt (14.5 ± 6.2), 17 nt (15.6 ± 6.0), and 19 nt (13.9 ± 5.1), while the 19 nt (20.6 ± 9.8) has moderately higher intensity in Cycle 3 and Cycle 4 than 11 nt (16.9 ± 8.1), 13 nt (17.2 ± 9.2), 15 nt (19.5 ± 9.8), and 17 nt (19.6 ± 10.0) (Supplementary Figure S1e). Overall, we determined that 15 nt is an appropriate length for the anchor primer. Finally, we also optimized the concentrations of interrogation probes and found that 0.2 μM (17.6 ± 7.2) is superior to 0.05 μM (11.5 ± 3.5) and 0.1 μM (12.9 ± 3.7) (Supplementary Figure S1f).

### IISS shows better specificity

Previously we found that false signals were occasionally generated when using mcPAL, leading to false decoding results. One of these examples was the sequencing of NEAT1 probe that mcPAL showed a high level of miscalling in the third position of the barcode, where an “A” was miscalled as a “G” due to fluorescence signals were present in two fluorescence channels (Figure 2a). However, only the correct Cy5 signal was observed when we changed to the 2-base encoding icPAL chemistry, showing better specificity of icPAL over mcPAL where the barcoding only depends on one single base (Figure 2c). CellProfiler were used to analyze the RCPs in the four sequencing cycles of these two SBL chemistry, showing that compared to RCPs in Cycle 1, the ratio of the false signals present in the Cycle 3 of the mcPAL is 58.19%, while only 1.15% for icPAL (Figure 2b and 2d). In order to avoid the influence of barcode sequence itself on the specificity of mcPAL, we re-designed the barcode of NEAT1 in mcPAL but false signals were still found in Cycle 3. For example, when the mcPAL barcodes of NEAT1 were changed to TTAA or GTTA, the false positive ratio of the Cycle 3 were still 76.54% and 55.63% respectively (Supplementary Figure S2a and S2b).

**Figure 2.**
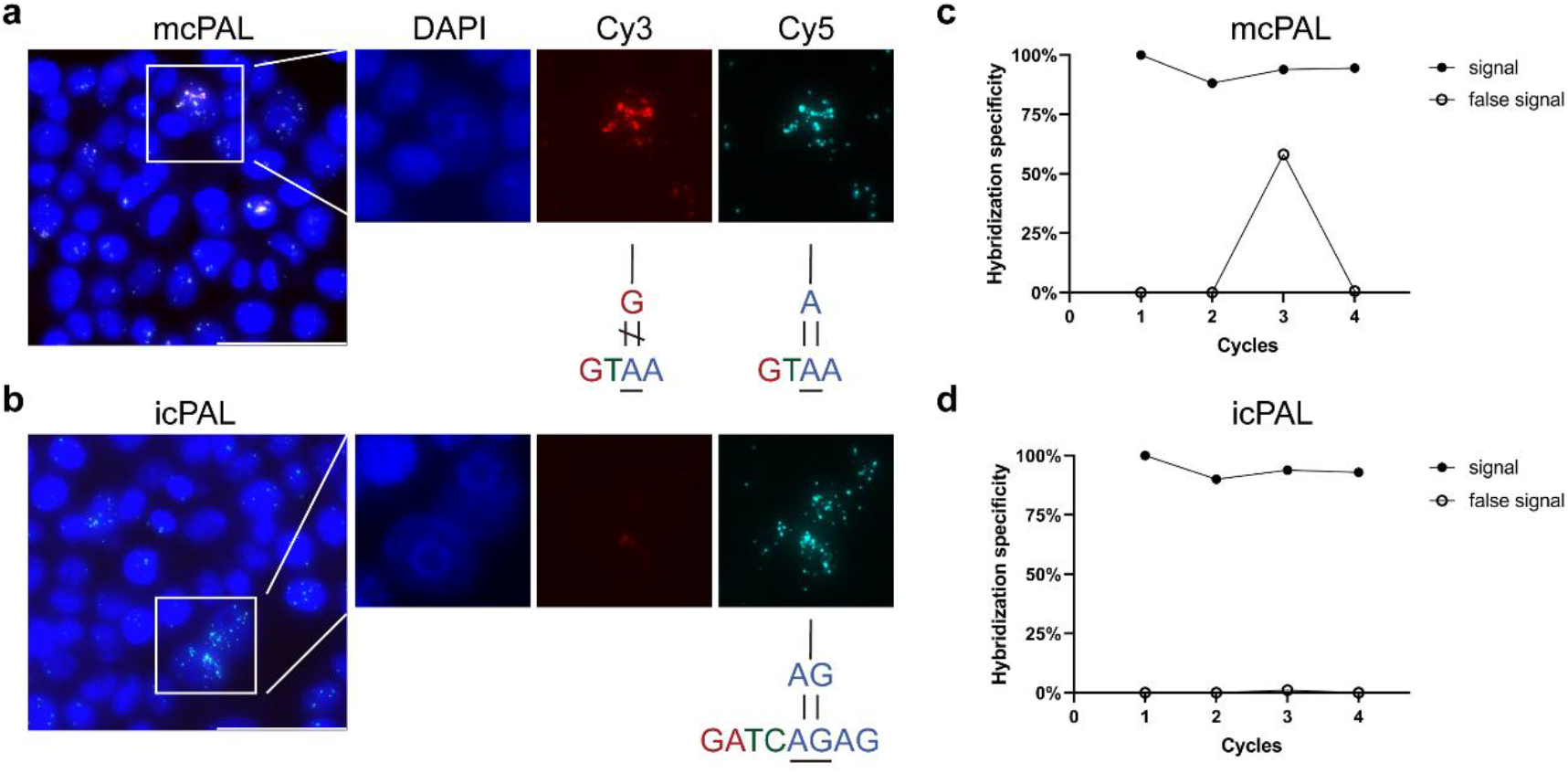
Comparison of icPAL-based IISS with mcPAL-based ISS. Detected RCP signal of the NEAT1 probe barcode in MCF-7 generated by the mcPAL-based chemistry (a) and the icPAL-based chemistry (b). Scale bar: 50 μm. (c, d) The ratio of the amount of correctly registered RCPs of NEAT1 in different cycles to that of Cycle 1 by mcPAL-based chemistry and icPAL-based chemistry.

In addition, we also compared the specificity between mcPAL and icPAL sequencing chemistries by detecting the barcode of another gene MALAT1 on MCF7 cell samples. We found that the mcPAL also had higher false signals in the Cycle 3 during *in situ* sequencing (60.87% for mcPAL v.s. 0.18% for icPAL) (Supplementary Figure S2c, d). These results showed that icPAL outperforms mcPAL in terms of specificity, thus can avoid false gene identification for *in situ* sequencing.

### IISS for *in situ* cell typing on fresh frozen mouse brain

To demonstrate the feasibility of IISS for *in situ* cell typing, we first performed IISS on a fresh frozen mouse brain tissue section (10 μm thick) (Supplementary Figure S3a). A panel of probes targeting 36 genes were curated based on previous publications ^17 18^, which allowed us to distinguish between main neuronal and non-neuronal cell types associated with mouse cortical and hippocampal region. To achieve better detection efficiency for single cells, three to five pairs of DLPs were designed for each gene. Three IISS cycles were required to decode these 36 genes (Supplementary Figure S3b). After decoding, we generated the spatial RNA expression atlas of these 36 genes in a mouse brain section (Supplementary Figure S3c). Specific spatial distribution patterns were observed for these genes (Supplementary Figure S3d). Cellpose was then used to segment cells based on nuclear DAPI staining so that the genes were assigned to individual cells, generating a gene-by-cell spatial gene expression matrix. Then we performed cell type annotation using a percentage-dominant methodology ^13^, in which each cell was annotated as a certain type that had the highest expression of respective marker gene panel. Results showed that the number of unique genes and RCPs made slightly difference among each cell types, fluctuating around 10 RCPs and 5 genes per cell, respectively (Figure 3a). Each marker panel was exclusively expressed in their corresponding cell types (Figure 3b). Finally, we defined seven types of cells (Figure 3c), including two types of neuronal cells (inhibitory neurons and excitatory neurons) and five types of non-neuronal cells (astrocytes, oligodendrocytes, vasculature, ependymal, and immune cells), which showed distinct spatial distributions on the expression map (Figure 3d).

**Figure 3.**
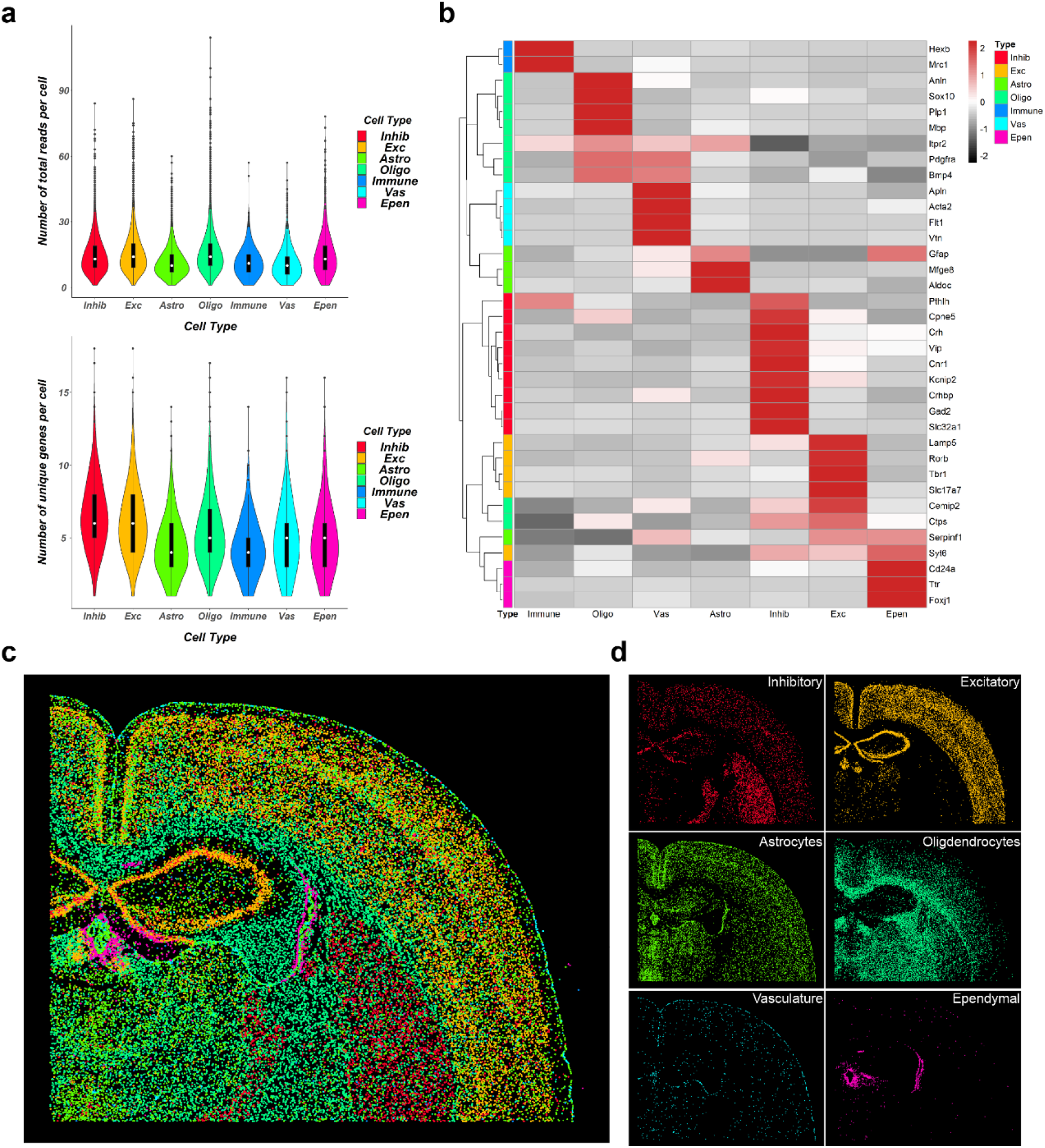
IISS for *in situ* cell typing on a fresh frozen coronal mouse brain tissue section. (a) Quality control violin plot, classified by cell type. The upper figure shows the total reads per cell according to different cell types, and the lower figure shows the number of genes per cell according to different cell types. (b) Gene expression heat map, where columns refer to cell types, and rows refer to genes. The rows have been hierarchically clustered. Red refers to high-expression level, while dark refers to low-expression level. (c) Overall cell type atlas and (d) single cell type atlas.

### IISS for cell type annotation, spatial trajectories inference and cell-cell interactions analyses in FFPE sample

Due to long preservation time and simple preservation conditions, archival FFPE tissue section is the major form for sample preparation both in clinical diagnosis and biological research. However, it may be more challenging to obtain *in situ* transcriptomic information from FFPE sections because of RNA degeneration and molecule cross-linking resulted from fixations. Thus, we established a working pipeline to perform *in situ* sequencing on FFPE sections. A spatial expression atlas of 40 genes was generated on an ulcerative moderately differentiated adenocarcinoma colon cancer section. These 40 genes are divided into four types, stromal cell, epithelial cell, immune cell and cancer-related signal pathway genes. Among them, immune cells are composed of B cells, T cells, macrophages, and mast cells. The spatial distribution of these 40 genes is shown in Figure 4a, while the necrosis regions have almost no RNA detected. According to the expression patterns of these genes, we divided this colon cancer section into stromal region, cancer region and cancer marginal area, which can be verified by comparing with hematoxylin and eosin (H&E) staining (Supplementary Figure S4a). And we observed the landscape of the tumor microenvironment of colon cancer through some marker genes of immune cells. For example, we have detected some macrophages with CD68 as a marker gene (Supplementary Figure S4b).

**Figure 4.**
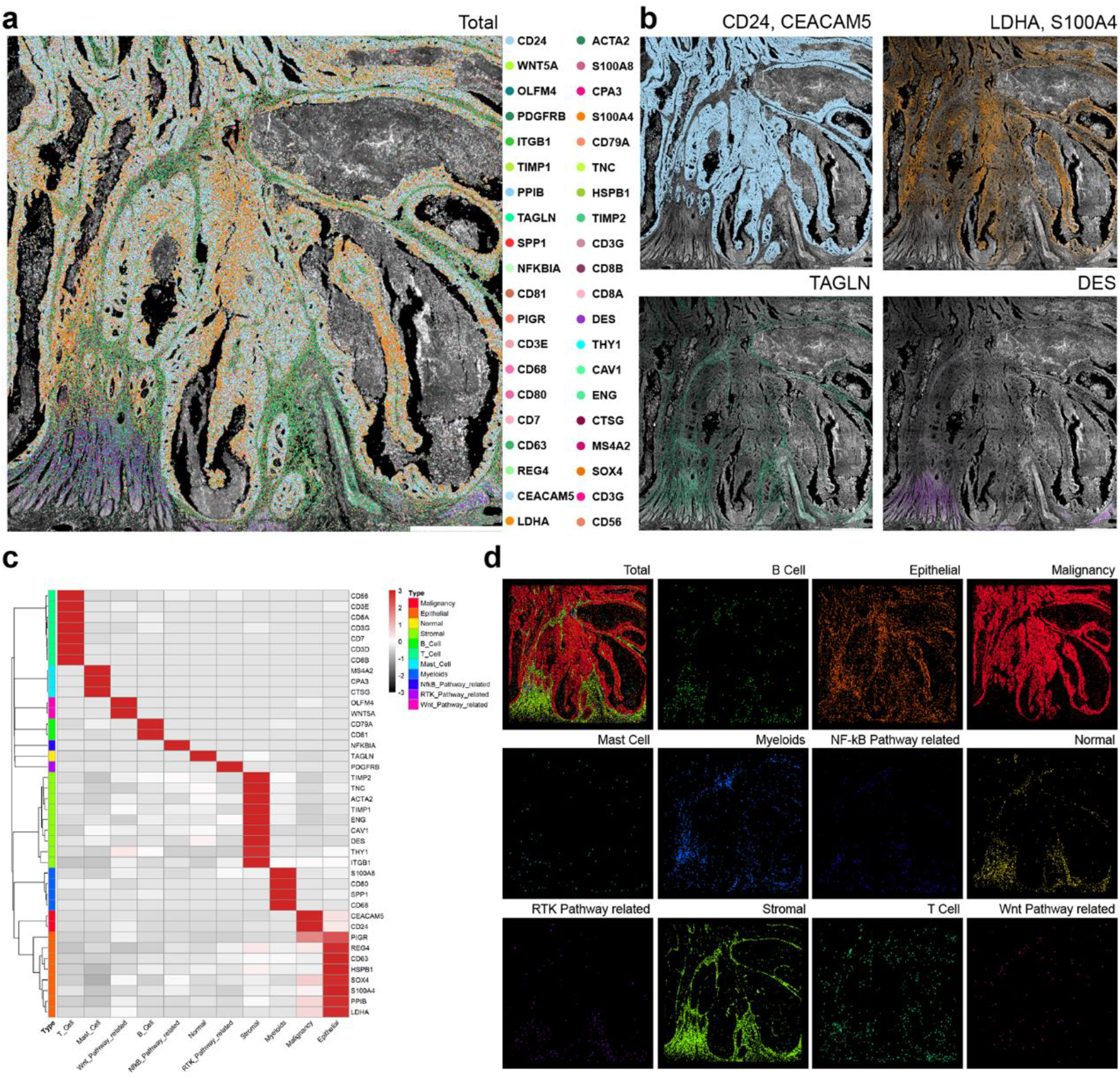
Single cell analysis of IISS on a colon cancer FFPE tissue section. (A) Overview of the 40-gene expression atlas in a colon cancer tissue section. Each color dot corresponds to a detected transcript signal. Scale bar: 10 mm. (b) Expression patterns of specific genes. CD24 and CEACAM5 colocalize in cancer regions. LDH and S100A4 colocalize in the cancer spreading region. TAGLN is located in matrix area. DES is mainly expressed in muscularis region. Scale bar: 10 mm. (c) Heat map of the expression patterns of different cell types. Columns are cell types, rows are genes, and the rows have been hierarchically clustered. (d) The spatial distributions of different cell types.

Apart from locating the expression distribution of individual genes, *in situ* single cell analysis can also be performed. Marker gene heatmap was firstly produced (Figure 4b), according to which can we see that the genes selected in this experiment are very representative and can be used as marker genes to represent the cell populations. In addition, we divided the cells on this colon cancer section into three categories by percentage-dominance cell typing methodology, namely histiocytes, immune cells and cells with high expression of cancer metabolizing pathway related genes. The histiocytes include epithelial cells, stromal cells, malignant cancer cells and normal cells. The immune cells include B cells, T cells, myeloid cells, and mast cells. The cancer signaling pathways include Wnt signaling pathway, NF-kB signaling pathway, and RTK signaling pathway. Then we visualized these cell types on the spatial map where a region-specific distribution pattern was observed (Figure 4c and 4d).

To understand the dynamic procedure of gene expression during the invasion of malignant tumor cells into stromal cells, and to find some key genes that may play an important role in treatment, we conducted a spatial trajectory inference analysis of target cell subsets of malignant tumor cells and stromal cells (Figure 5a). The evolutionary trajectories are reconstructed based on transcriptome profiles and the spatial context of cells within the tissue. We also performed trajectory-based differential expression analysis and selected two representative branches (branch 6318 and 7182) to observe gene changes in the trajectory development (Figure 5b). Along the development of the trajectory, the expression of T cell and mast cell genes tended to decrease, while DES, CAV1 and THY1 of cancer-related fibroblasts increased, indicating that cancer-related fibroblasts overexpressed by these genes may play an important role in promoting cancer metastasis and invasion ^19,20^. To explore the possible immune process, we also performed trajectory analysis inference from myeloids to stromal cells and analyzed expression changes of genes in key branches (Figure 5a and 5b). In clade 9874, some genes were down-regulated along the trajectory, such as NFKBIA, whose expression was associated with the ability of colon cancer cells to infiltrate and metastasize. It is an NF-κB pathway-related gene, and it has been shown that the NF-κB pathway activates dendritic cells to drive anti-tumor immunity ^21^.

**Figure 5.**
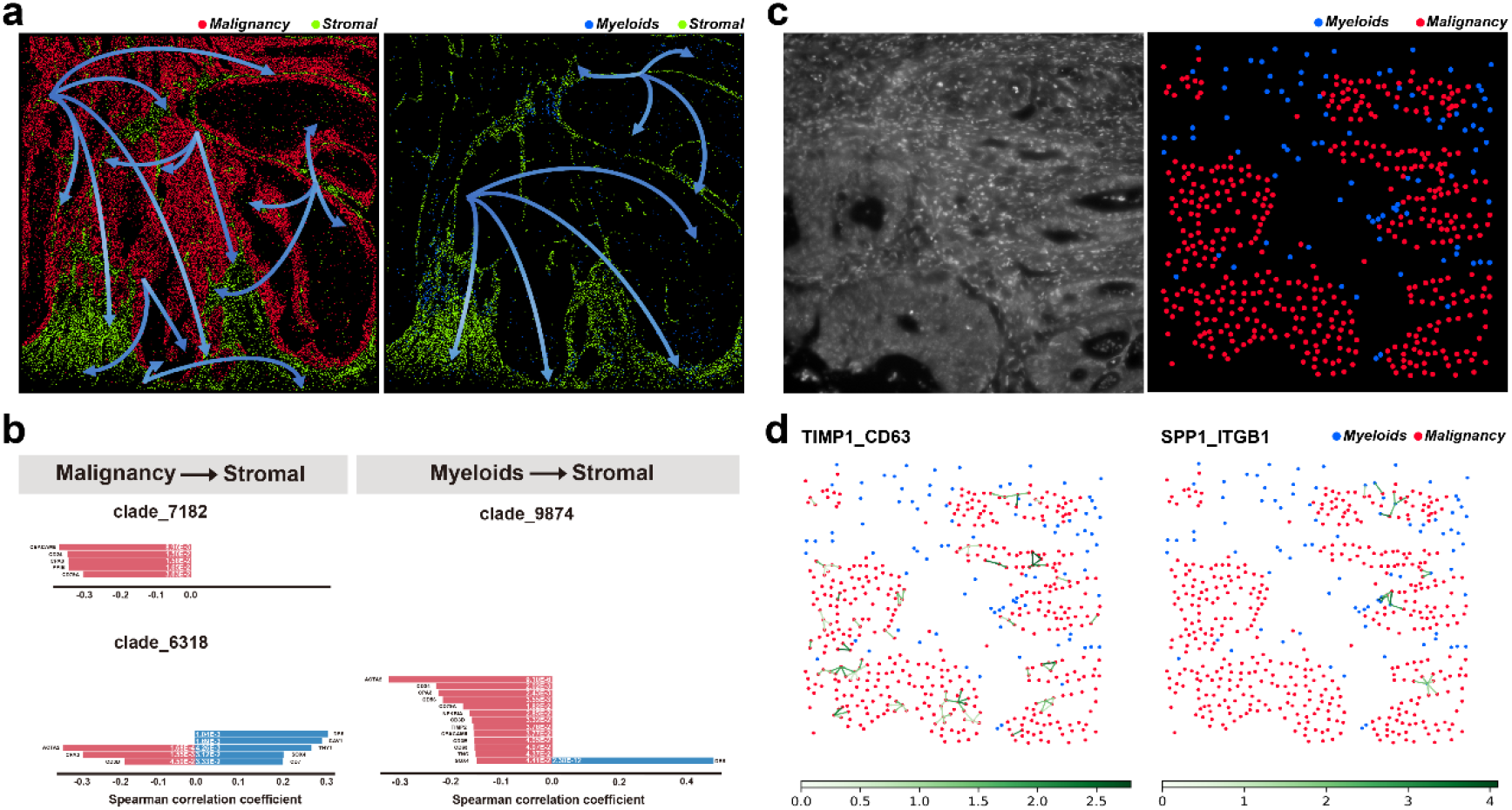
Spatial trajectory inference and cell-cell interaction analysis. (a) Developmental trajectories of malignancy towards stromal, and myeloids towards stromal. (b) Trajectory-related genes. Genes from the left side (red) are negatively correlated with spatial trajectory, while genes from the right side (blue) are positively correlated. (c) Cell type annotation results for the target field. (d) Visualization of ligand-receptor interactions between different cell types on a featured FOV

Finally, we analyzed cell-cell interactions (CCI) on this tissue by combining cell-type diversity and L-R expression information to identify regions with the highest likelihood of interaction ^22,23^. Using the LR database available in stlearn, five pairs of LRs were identified from our 40 genes with P-value < 0.05, and they were ranked based on significance level. We selected the top LR (CD63-TIMP) and a pair of LR (SPP1-ITGB1) associated with tumor metastasis from these five pairs of LRs to perform significant CCIs prediction (Supplementary Figure S4d). We chose a featured FOV with the most obvious interaction between cells and visualized the distribution of target cell types which included malignancy and myeloids (Figure 5c and 5d). Previous study showed that patients with high expression of SPP1 have a poor prognosis ^24^. According to our analysis, SPP1 gene expression is related to ITGB1, and also to some cancer-related fibroblast subsets. Cancer related fibroblasts secrete various cytokines, growth factors, chemokines, exosomes and other effector molecules to interact with tumor infiltrating immune cells and other immune components in TME, thus forming immunosuppressive TME and allowing cancer cells to escape the immune surveillance system ^25^. In addition, SPP1 overexpressing macrophages are located at the junction of cancer and stromal areas, which may have a great relationship with the metastasis and infiltration of tumors. In respect to another LR pair, CD63 is a surface receptor of exosomes ^26^, and TIMP1 may act as a signal molecule in this process.

## DISCUSSION

As a further development of the original *in situ* sequencing technology, the IISS method is based on a modified probe design scheme combined with novel barcoding strategy using combinatorial 2-base encoding to improve the sequencing quality. Although we also tested combinatorial 3-base encoding scheme, the results still showed that the 2-base encoding was better than the 3-base encoding scheme (Supplementary Figure S5a and S5b). Previously, we demonstrated that the DLPs have better detection efficiency and higher specificity than the padlock probes in the RNA-templated *in situ* hybridization assay ^15^. Here, the *in situ* sequencing library for targeted spatial transcriptomic gene expression analysis constructed by DLPs can also benefit from their advantages. While the use of split barcode on the upstream and downstream DLP pair may also help to exclude false ligations, because only correct barcodes called by IISS will be picked up. We also optimized the *in situ* sequencing protocol and determined the optimal probe concentration as well as the anchor probe length. More importantly, we found that one-step sequencing where the hybridization of anchor primer, interrogation probe and ligation was performed simultaneously resulted in similar quality as that of the two-step sequencing protocol where anchor primer is hybridized first and interrogation probe is ligated in a separate step. This helped to simplify the sequencing protocol and save reaction time. A glitch in the current IISS protocol is the high detection efficiency can sometimes cause signal overcrowding due to regionally high expression of certain genes. This can result in loss of gene expression information in those regions because of signal registration failure due to the image analysis pipeline only recognizes dot-like signal but the oversized signal plaques are filtered out (Supplementary Figure S5c and S5d). By examining the raw images, we noticed that a large proportion of RCPs encoding Ttr are missing due to this reason.

A key step in the spatial transcriptomics analysis is to generate single cell gene expression matrix *in situ*. Thus, in the IISS-based targeted spatial transcriptomic gene expression analysis pipeline, we established an analysis pipeline that exploited Cellpose for cell segmentation combined with percentage-dominance cell typing. Based upon the *in situ* cell atlas generated by the pipeline, advanced single cell analysis, such as spatial trajectories inference and cell-cell interactions, can also be performed. Thus, benefiting from these enhancements, we can not only compute a spatial distribution atlas of neuronal and non-neuronal cells on fresh frozen mouse brain tissue, but also on cancer FFPE tissue samples. We surprisingly found that the cancer-related fibroblasts overexpressing DES, CAV1 and THY1 may contribute to the carcinogenesis from normal region towards malignancy region, showing its potentials to help decipher novel gene functions. Previously, most of the imaging based spatial transcriptomic methods were performed on fresh frozen tissue sections. Here, we established the experimental pipeline and demonstrated that advanced spatial gene expression analysis can also be perform on FFPE tissue sections, showing its great potential for clinical diagnosis.

In summary, we have established a novel full pipeline for high-resolution targeted *in situ* gene expression profiling. The high specificity and detection efficiency of IISS make it an ideal imaging-based technique that can be easily implemented and adapted to various tissue types.

## MATERIAL AND METHODS

### Gene selection and probe design

The genes used for icPAL evaluation on cell samples were randomly selected from our in-house probe storage, while the mouse brain gene panel and the genes selected for colon cancer tissue were manually selected based on existing literatures ^17,18,24,27-29^. Probes were designed using asmFISH DNA ligation probe (DLP) designing principle ^15^, thus that the barcodes were divided into two motifs placed in the upstream and downstream probes that constitute the DLP pairs respectively (Supplementary Fig. S1). The length of each probe hybridized to RNA target is 16 nt, making the sum of the whole target template 32 nt long for each DLP hybridization site. The probe hybridization targets were selected by using BLAST (https://blast.ncbi.nlm.nih.gov/Blast.cgi) to ensure their specificity. Three to five targets were selected per gene to ensure the detection efficiency. DLPs were phosphorylated in a mix containing 0.2 U/μL T4 polynucleotide kinase (Thermo Fisher Scientific), 1× PNK buffer A (Thermo Fisher Scientific), 1 mM ATP (Thermo Fisher Scientific), 2 μM DLPs, and incubated at 37°C for 30 min and 65°C for 10min. The interrogation probes were ordered from Invitrogen (Shanghai, China), while all the rest of oligonucleotides were ordered from Sangon (Shanghai, China). The sequence information is listed in Supplementary Table S1.

### Cells

MCF-7 cells (ATCC) were cultured in DMEM 1640 (Cat No. L140KJ; BasalMedia, Shanghai, China) supplemented with 10% fetal bovine serum (FBS, Cat No. A0500-3011; Cegrogen Biotech, Germany) for 24-48 h. Then the cells were treated with trypsin (Cat No. D121001; BasalMedia, Shanghai, China) to suspension and seeded on sterile slides (Cat No. 80312-3161; CITOTEST, Jiangsu, China). Cells were allowed to grow and adhere on the slides for 12-24 h, followed by washing twice with 0.1% diethyl pyrocarbonate (Cat No. D5758; Sigma-Aldrich, MO, USA) treated with 1× phosphate buffered saline (PBS, Cat No. G211014; BasalMedia, Shanghai, China) (1× DEPC-PBS). Fixation was performed using 4% paraformaldehyde (PFA, Cat No. 16005; Sigma-Aldrich) in 1× DEPC-PBS for 30 min at room temperature. Next, the slides were washed with 1× DEPC-PBS twice, dehydrated with an ethanol series of 70%, 85%, and absolute for 5 min each, and then stored at −80°C until used. The slides were warmed up to room temperature and dried before used.

### Tissue samples

All experiments were performed in accordance with relevant guidelines and regulations. C57BL/6 mice were ordered from SLAC Laboratory Animal (Shanghai, China). Mouse brain was embedded and frozen in optimal cut temperature (OCT) (SAKURA, USA) compound at - 80°C and 10 μm cryosection slices were prepared by using a Leica cryostat (Leica, Germany). The slices were warmed up to room temperature and fixed with 4% PFA for 5 min before used. After washing twice with 1× DEPC-PBS for 2 min each, dehydration was performed in ethanol series of 70%, 85%, and absolute for 1 min each and washed three times with 1× DEPC-PBST (1x DEPC supplemented with 0.05% Tween-20, Cat No. P9416; Sigma-Aldrich, MO, USA). Permeabilization of tissue was carried out in 0.1 M HCl for 5 min and then washed three times with 1× DEPC-PBST.

Formalin-fixed, paraffin-embedded (FFPE) tissue sections of colon cancer were obtained from the Pathology Department of the 910 Hospital, Quanzhou, Fujian, China. Deparaffinization was firstly performed on FFPE tissue section slides which were baked at 60°C for 30 min and then dipped into xylene twice at room temperature for 15 min and 10 min respectively, followed by submerging in absolute ethanol twice, 95% ethanol twice, and 70% ethanol twice for 2 min each time, and then washed with DEPC-H_2_O for 5 min and with 1× DEPC-PBS for 2 min. Fixation was performed in 4% PFA in 1× DEPC-PBS for 10 min and washed twice with 1× DEPC-PBS. Before permeabilization, 30 mL of 0.1 M HCl (Cat No. 10011018; SINOPHARM, China) was warmed up to 37°C for subsequent pepsin penetration. Permeabilization was carried out by submerging the slides in 0.1 M HCl added with 30 μL of 100 mg/mL pepsin (Cat No. P7102; Sigma-Aldrich) for 30 min at 37°C. Slides were then washed with DEPC-H_2_O for 5 min and with 1× DEPC-PBS for 2 min, incubated in ethanol series of 70%, 85%, and absolute for 1 min each for dehydration, finally air dried and washed three times with 1× DEPC-PBST.

The use of animal tissues and tumor tissue in this study was approved by the ethics committee of School of Medicine, Huaqiao University, Quanzhou, Fujian, China.

### DLP hybridization and ligation

ImmEdge hydrophobic barrier pen (VECTOR LABS, Cat No. H-4000; CA, USA) was used to create a reaction area. To hybridize the DLPs with target mRNA, 0.1 μM of phosphorylated DLPs in 6× SSC and 10% formamide was added on the slides and incubated for 4 h at 37°C. After washing three times with 1× DEPC-PBST, the upstream and downstream probes of DLP pairs was ligated by applying a ligation mix containing 1× SplintR buffer (Cat No. B0375S; NEB, Beijing, China), 2.5 U/μL SplintR Ligase (Cat No. M0375 L, NEB, Beijing, China), 1 U/μL RiboLock RNase Inhibitor (Cat No. EO0384, Thermo Fisher Scientific), 25% glycerol (Cat No. G5516; Sigma-Aldrich), and 0.2 μg/μL BSA (Cat No. B2064; Sigma-Aldrich), and incubated for 0.5 h at 37°C. Then the slide was washed three times with 1× DEPC-PBST.

### DLP circularization and RCA

The ligated upstream and downstream probes of DLP pairs were circularized by DNA ligation assisted by a splint primer (Supplementary Table S1). First, 0.5 μM of splint primer in 6× SSC and 10% formamide was added to the reaction area and incubated for 0.5 h at 37°C, allowing hybridization with the ends of the DLPs. Probe circularization and RCA was initiated by applying a reaction mix in DEPC-H_2_O containing 1× Phi29 buffer (Cat No. EP0094; Thermo Fisher Scientific), 1 U/μL Phi29 DNA polymerase (Cat No. EP0094; Thermo Fisher Scientific), 0.1 U/μL T4 DNA ligase (Cat No. EL0011; Thermo Fisher Scientific), 1 mM dNTP (Cat No. R0182; Thermo Fisher Scientific), 5% glycerol, 0.2 μg/μL BSA, and 1 mM ATP to the reaction area and incubated overnight at room temperature, then washed three times with 1× DEPC-PBST.

### Decoding of RCPs by *in situ* sequencing

For sequencing, a mix containing 200 nM anchor primer (Sangon Biotech, Shanghai, China), 1× T4 DNA ligase buffer (Cat No. EL0011; Thermo Fisher Scientific), 0.1 U/μL T4 DNA ligase, 200 nM fluorescently labeled sequencing interrogation probes (Invitrogen, Shanghai, China), 2 mM ATP, and 0.2 μg/μL BSA were applied and incubated at 30°C for 45 min, followed by washing three times with 1× DEPC-PBST. The slides were then mounted in SlowFade Gold Antifade Mountant (Fermantas) medium containing 0.5 μg/mL DAPI. Then the images were acquired by a Leica DM6B fluorescence microscope (Leica, Germany) equipped with a DFC9000GT camera and a DFC7000T camera (Leica, Germany) using a 20× objective (Leica, Germany). To strip off ligated sequencing probes and prepare the slides for the next sequencing cycle, the slides were treated six times with stripping buffer that contained 80% formamide and 0.1% Triton X-100 at 37°C for 10 min each and washed three times with PBST. The same procedures were applied for each sequencing cycle by repeating the hybridization and ligation of sequencing probe, imaging and stripping off.

### Image analysis

For image registration and base calling, Z-stacks and maximal intensity projection (MIP) were applied in all imaging processes. For cell samples, MIP images were analyzed with the open-source image analysis software CellProfiler (version 4.0.3) (https://cellprofiler.org/) to quantitatively measure the integrated fluorescence intensity of rolling circle amplification products (RCPs). All images were exported as single channel .tif files subjected for CellProfiler analysis. The module “*MeasureObjectIntensity*” was used to obtain the fluorescent intensity of RCPs. For both fresh frozen tissue sections and FFPE tissue sections, image registration was firstly carried out for each field of view (FOV) by image correction on multiple cycles, then aligning them with the reference stitched image generate by Leica Application Suite X Life Science (Leica, Germany) with 10%. Finally, the relationship between gene and RCPs can be determined by decoding their color encoding schemes.

### *In situ* cell typing analysis

To distinguish among different cell types in single cell level, we performed cell segmentation and cell type annotation. For cell segmentation, we used Cellpose ^16^ to identify cell nuclei on DAPI staining images. Specifically, featured FOV were selected manually and segmented using built-in “*Cyto*” model. Then, a new model was trained on each FOV, which was then used to segment the full view. A .npy file was obtained as an output which stores all necessary information about cell segmentation. To include the gene information distributed on cytoplasm, the identified nuclei were then expanded 20 pixels to represent cell area. Finally, each fluorescent signal was assigned to cell using a customized python script. A gene-by-cell matrix was generated and used in all downstream analyses. A cell type scoring methodology resembling that of pciSeq ^13^ was exploited to classify cell class. For each cell, the value of raw expression matrix was converted to percentage value, so the sum of expression value is equal to 1. Then, we calculated as scores the sum of probability of each cell type-related marker gene panel. A cell was labelled as a certain class with the highest score. Finally, we filtered out the ambiguous cells that can be annotated as arbitrary cell types.

### Spatial trajectory inference and cell-cell interactions

Both spatial trajectory inference and cell-cell interactions are based on stLearn ^22^, which was exploited to perform spatial trajectory reconstruction. We firstly input raw expression matrix, cell centroid locations and cell type labels to create an AnnData object. Outliers were excluded using “*filter_cells*” with “*min_genes = 1*” and “*min_counts = 5*”. Then, a standard data pre-processing process, including log normalization, scaling and PCA, was performed on filtered data.

From the beginning of spatial trajectory inference pipeline, we chose the cells labelled with malignancy as root. Then we ran the global level of pseudo-time-space (PSTS) method to reconstruct the spatial trajectory between malignancy and stromal cells. Based on the spatial trajectory, we can see certain clades are started from sub-malignancy and sub-stromal. Then we ran “*spatial*.*trajectory”*, “*detect_transition_markers_clades”* to detect the highly correlated genes with the PSTS values. The same method was used for analyzing the spatial trajectory between myeloids and stromal cells.

stLearn ^22^ and the same data pre-processing process as mentioned in above section were used in deciphering cell-cell interactions (CCIs), except logarithmizing and scaling. The first step in the stLearn CCI pipeline was to perform both ligand-receptor (LR) analysis and p-value adjustment to find significant LR pairs. With the establishment of significant areas of LR interactions, we predicted significant CCIs, computed CCI network and plotted spatial cell type interactions using “*tl*.*cci*.*run_cci”* and “*pl*.*lr_plot”*, respectively.

## DATA AVAILABILITY

Original data of the *in situ* sequencing is available on reasonable request to the corresponding authors.

## CODE AVAILABILITY

The custom codes used to analyze the IISS data are deposited on Github (https://github.com/HQU-MDLAB/IISS).

## ACKNOWLEDGEMENTS

We thank Prof. Diao and Prof. Lin for valuable discussions on this project. This study was supported by the Natural Science Foundation of Fujian Province (2022J06022), Quanzhou Science and Technology Plan Project (2021C040R) and the Scientific Research Funds of Huaqiao University.

