## Supplementary Figures for "Improved *in situ* sequencing for high-resolution targeted spatial transcriptomic analysis in tissue sections"

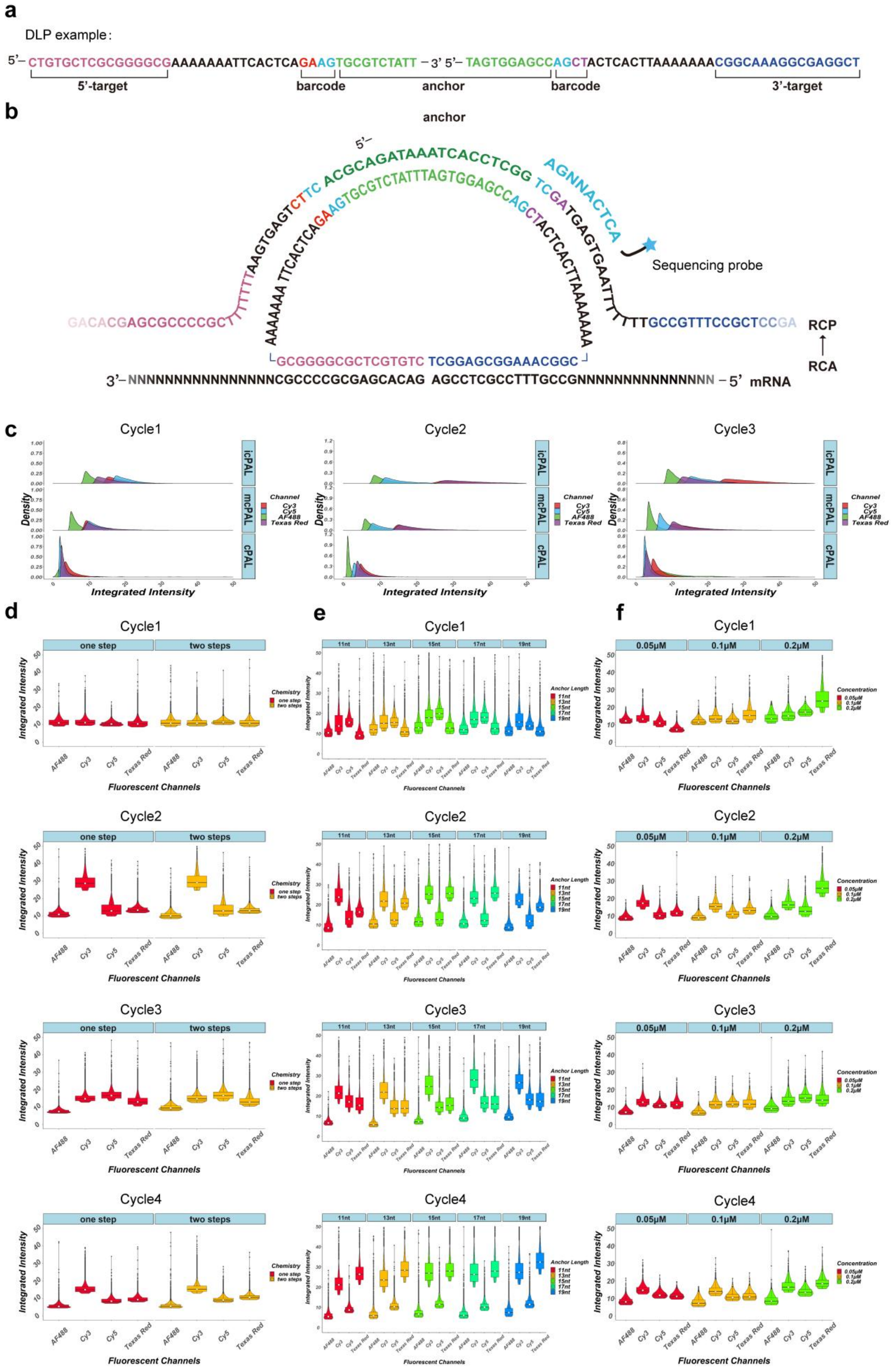


Supplementary figure S1. Overview of IISS probe design and comparison of integrated intensity among different experimental conditions using icPAL sequencing chemistry. **(a)**

Schematic representation of a DLP pair. Each pair is constituted by two probes. Left probe contains a 16 nt target recognition sequence, 10 nt unrelated sequence, a 9 nt interrogation probe sequence and a 10 nt anchor primer hybridization sequence from 5' to 3', assembling a 45 nt long DNA sequence, while all the elements are reversely distributed in right probe from 5' to 3'. **(b)** DLP hybridization, circularization, RCA, and decoding of RCPs. **(c)** Comparison of maximum integrated intensity of fluorescence signal in different cycles among different sequencing chemistries, namely icPAL, mcPAL, and cPAL. **(d)** Comparison of maximum integrated intensity of fluorescence signal in different cycles between one-step and two-steps sequencing protocols. **(e)** Comparison of maximum integrated intensity of fluorescence signal in different cycles between different anchor length, namely 11 nt, 13 nt, 15 nt, 17 nt, and 19 nt. **(f)** Comparison of maximum integrated intensity of fluorescence signal in different cycles between different concentrations of interrogation probes, namely 0.05 μM, 0.1 μM, and 0.2 μM.

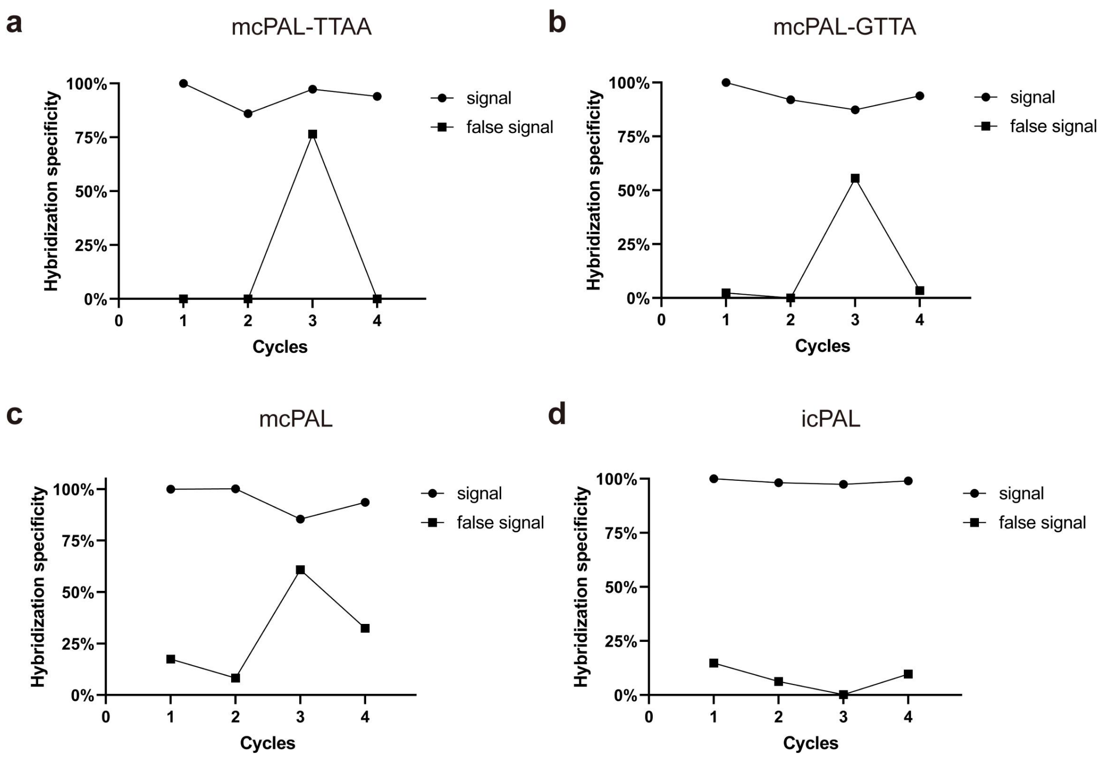


Supplementary Figure S2. Comparison of hybridization specificity of mcPAL-based ISS with different barcode designed, and between mcPAL- and icPAL-based ISS. **(a)** The ratio of the amount of correct registration RCPs of NEAT1 in 4 Cycles to that of Cycle 1 using mcPAL-based chemistry. Barcode: TTAA. **(b)** The ratio of the amount of correct registration RCPs of NEAT1 in 4 Cycles to that of Cycle 1 using mcPAL-based chemistry. Barcode: GTTA. **(c)** The ratio of the amount of correct registration RCPs of MALAT1 in 4 cycles to that of the Cycle 1 using mcPAL-based chemistry. **(d)** The ratio of the amount of correct registration RCPs of MALAT1 in 4 cycles to that of the Cycle 1 using icPAL-based chemistry.

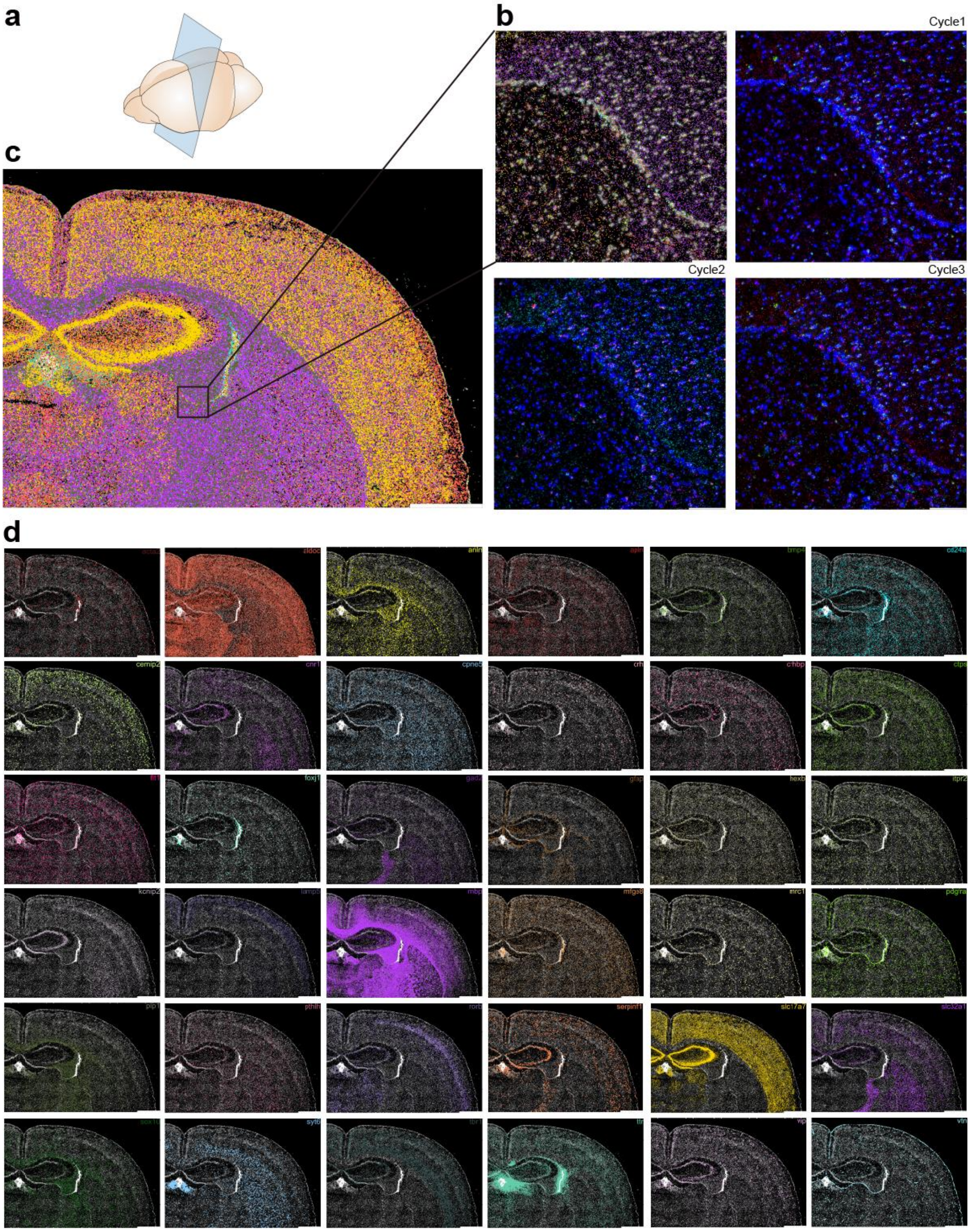


Supplementary Figure S3. IISS on fresh frozen mouse brain. **(A)** Illustration of tissue section used from whole mouse coronal sections used for IISS. **(B)** Representative FOVs. Scale bar: 100 µm. **(C)** Decoding of 36 genes over 3 rounds of *in situ* sequencing. Each dot is a transcript detected, and each color corresponds to a unique gene. Scale bar: 10mm. **(D)** Spatial distribution of single genes. Scale bar: 10mm.

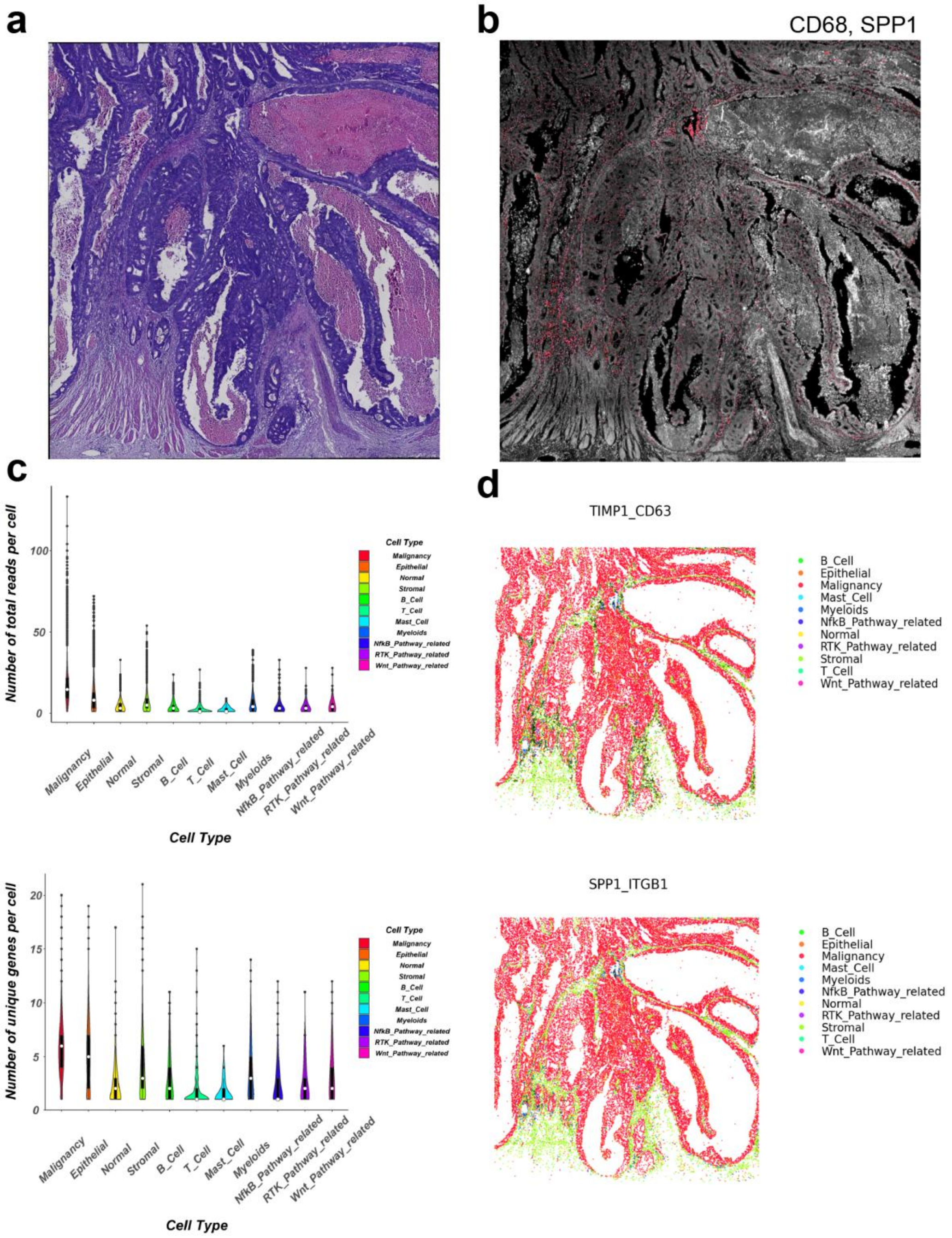


Supplementary Figure S4. H&E image of colon cancer FFPE tissue and corresponding spatial analysis results. **(A)** H&E image shows this section contains cancer region, matrix region, and necrosis region. **(B)** IISS results showing the distribution of SPP1$^{+}$ macrophages with CD68. **(C)** Statistical data of IISS results at a per cell basis. The upper figure shows the total reads per cell, and the lower figure shows the number of genes per cell. **(D)** Ligand-receptor interactions between different cell types on the whole tissue section. TIMP1 and SPP1 are ligands targeting CD63 and ITGB1, respectively.

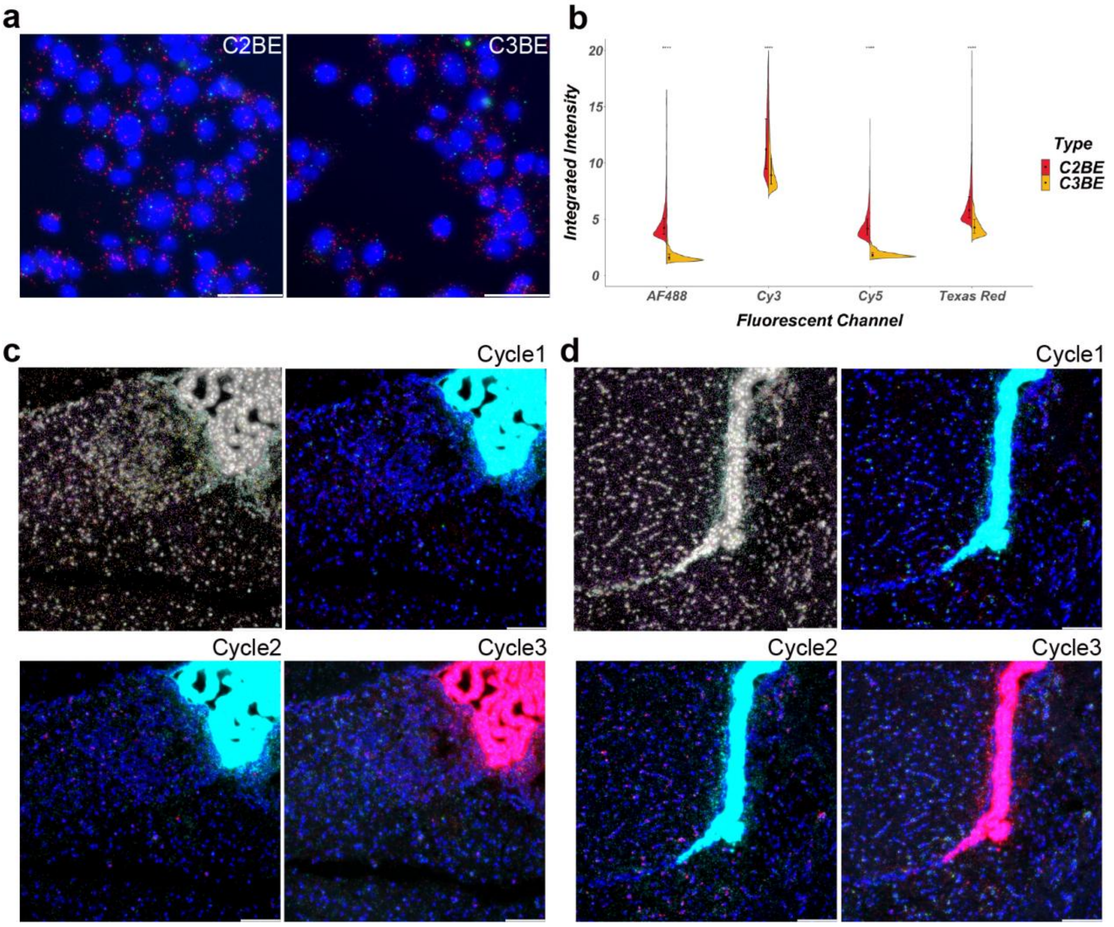


Supplementary Figure S5. Comparison between combinatorial 2-base encoding v.s. combinatorial 3-base encoding and optical crowding on mouse brain section. **(a)** Implementation of combinatorial 2-base encoding (C2BE) and combinatorial 3-base encoding (C3BE) ISS on MCF-7 cell slides. **(b)** Comparison of maximum integrated intensity of fluorescence signal between C2BE and C3BE. **(c)** and **(d)** Optical crowding of Ttr signal on featured FOVs among three cycles of sequencing.
